# Immortalised hippocampal astrocytes from 3xTG-AD mice fail to support BBB integrity *in vitro*: Role of extracellular vesicles in glial-endothelial communication

**DOI:** 10.1101/2020.03.26.009563

**Authors:** Karolina Kriaučiūnaitė, Aida Kaušylė, Justina Pajarskienė, Virginijus Tunaitis, Dmitry Lim, Alexei Verkhratsky, Augustas Pivoriūnas

## Abstract

Impairments of the blood brain barrier (BBB) and vascular dysfunction contribute to Alzheimer’s disease (AD) from the earliest stages. However, the effects of AD-affected astrocytes on the BBB remain largely unexplored. In the present study we created an in vitro BBB using human immortalised endothelial cells in combination with immortalised astroglial cell lines from the hippocampus of 3xTG-AD and wild-type mice (3Tg-iAstro and WT-iAstro, respectively). We found that co-culturing endothelial monolayers with WT-iAstro up-regulates expression of endothelial tight junction proteins (claudin-5, occludin, ZO-1) and increases the trans-endothelial electrical resistance (TEER). In contrast, co-culturing with 3Tg-iAstro does not affect expression of tight junction proteins and does not change the TEER of endothelial monolayers. The same in vitro model has been used to evaluate the effects of extracellular vesicles (EVs) derived from the WT-iAstro and 3Tg-iAstro. The EVs derived from WT-iAstro increased TEER and up-regulated expression of tight junction proteins, whereas EVs from 3Tg-iAstro were ineffective. In conclusion, we show for the first time that immortalised hippocampal astrocytes from 3xTG-AD mice exhibit impaired capacity to support BBB integrity in vitro through paracrine mechanisms and may represent an important factor underlying vascular abnormalities during development of AD.

## Introduction

Impairments of the blood-brain barrier (BBB) contribute to pathological evolution of Alzheimer’s disease (AD) from its earliest stages. Functional imaging studies detected delayed cerebral blood flow response and reduction of glucose uptake in individuals with mild cognitive impairment (MCI), which is considered as a prodromal stage of AD (Rombouts et al. 2005; Hunt et al. 2007). Furthermore, advanced imaging studies demonstrated that BBB breakdown in the hippocampus preceded clinical manifestation of MCI (Montagne 2015). Loss of BBB function and integrity is manifested in both familial and sporadic AD; in particular multiple “vasculotropic” mutations in amyloid precursor protein cause breakdown of BBB and are associated with cerebral amyloid angiopathy (Basun et al. 2008; Grabowski et al. 2001; Zarranz et al. 2016); BBB integrity is also severely compromised in humans carrying presenilin mutations (Dermaut et al. 2001; Lemere et al. 1996; Szpak et al. 2007). Similarly, compromised BBB and amyloid angiopathies have been frequently documented in transgenic AD animals expressing mutant AD-related genes (see (Sweeney et al. 2019) for details and relevant references). Astrocytes play a fundamental role in the formation and maintenance of BBB (Sweeney et al. 2019; Alvarez et al. 2013; Srinivasan et al. 2015) and therefore astroglial dysfunction, which occurs at the early stages of the disease may be critical for BBB damage. Reduction of brain vascular volume and a thickening of collagen-laden basement membrane of the endothelium surrounding microvessels emerged in hippocampi of 3xTG-AD mice well before the appearance of senile plaques and neurofibrillary tangles (Bourasset et al. 2009; Do et al. 2014). Functional impairments of BBB are detected in AD transgenic models as early as at 1-3 month of age, while β-amyloid depositions and cognitive deficits develop later at 9 – 12 months of age (Paul et al. 2007; Sagare et al. 2013; Ujiie et al. 2003). Our previous studies on 3xTG-AD mouse model (Olabarria et al. 2010)^18^ as well as on stem cells derived human astrocytes (Jones et al. 2017) revealed astroglial atrophy developing in the pre-plaque stages of the disease, therefore asthenia of astrocytes accompanied with the loss of function may contribute to the early impairment of the BBB.

Several attempts have been recently made to construct an *in vitro* model of BBB using brain microvascular endothelial cells growing on membrane-based or extracellular matrix-based platforms (Katt et al. 2018; Bogorad et al. 2015; Campisi et al. 2018; Phan et al. 2017; Motallebnejad et al. 2019; Shin et al. 2019). In the present study we created an *in vitro* membrane-based chimeric BBB using human immortalised endothelial cells in combination with recently generated immortalised astroglial cell lines from the hippocampus of 3xTG-AD and wild-type control mice (3Tg-iAstro and WT-iAstro, respectively) (Rocchio et al. 2019). These lines represent useful experimental tools allowing generation of large numbers of cells with properties similar to primary hippocampal astrocytes. 3Tg-iAstro also show alterations in transcription and deregulation of Ca^2+^ signalling previously described in primary cultures derived from hippocampi of 3xTG-AD mice (Rocchio et al. 2019). Therefore we aimed at clarifyinmg whether these immortalised astrocytes carry the functional pathological signature associated with AD pathology; in particular we focused on the potential inability of diseased astrocytes to support endothelial monolayer barrier function.

In the present study we analysed how WT-iAstro and 3Tg-iAstro affect trans-endothelial electrical resistance (TEER) and expression of tight junction proteins when co-cultured with brain endothelial cell monolayers. We found that WT-iAstro up-regulate expression of tight junction proteins in endothelial monolayers, which in turn increases the TEER. In contrast, co-culturing with 3Tg-iAstro does not affect expression of tight junction proteins and does not change the TEER of endothelial monolayers. The same *in vitro* model has been used to evaluate the effects of extracellular vesicles (EVs) derived from the WT-iAstro and 3Tg-iAstro. The EVs derived from WT-iAstro increased TEER and up-regulated expression of tight junction proteins, whereas EVs from 3Tg-iAstro were ineffective. We also found that incubation with conditioned medium from brain endothelial cells increased secretion of the pro-inflammatry chemokine CCL2 from 3Tg-iAstro but not from WT cells. We contemplate the existence of bi-directional communications between endothelial cells and astrocytes needed for maintaining BBB. Astrocytes affected by AD pathology loose the ability to support the integrity of BBB through compromised paracrine communication. This further corroborates the idea of astroglial paralysis in the context of AD: asthenic astrocytes are incapable of maintaining BBB thus adding to pathological evolution of the disease.

## Results

### WT-iAstro, but not 3Tg-iAstro increase transendothelial electrical resistance (TEER) of endothelial monolayers

To create the chimeric model of the BBB *in vitro* we combined two cellular monolayers – the immortalized human brain endothelial cells (hCMEC/D3) were placed on the polyester membrane transwell inserts, whereas astrocytes (that represent the “brain” part of the BBB) were seeded at the bottom of the wells (Fig. 1a). The distance between two cellular layers was ∼ 1.5 mm which allowed paracrine communications. Endothelial cells growing on the membranes established intercellular contacts and expressed tight junction proteins, which secure the barrier. Monitoring electrical resistance of endothelial monolayers allows quantitative readout of the barrier integrity ^12^. The TEER of the endothelial monolayer cultured alone was 10.11 ± 0.717 W (n = 11). Co-culturing hCMEC/D3 cells with WT-iAstro for 6 days significantly increased the TEER of an *in vitro* BBB to 11.95 ± 0.44 W (or by 18.2 ± 4.41 %, n = 14, p < 0.05; Fig. 1b). In contrast, when endothelial cells were cultured with 3Tg-iAstro, the TEER has not been affected (TEER reading in the presence of 3Tg-iAstro was 9.99 ± 0.68 W, n = 14; Fig. 1b). Thus AD pathology suppresses the astroglial mechanisms supporting the integrity of endothelial barrier *in vitro*.

**Fig. 1.**
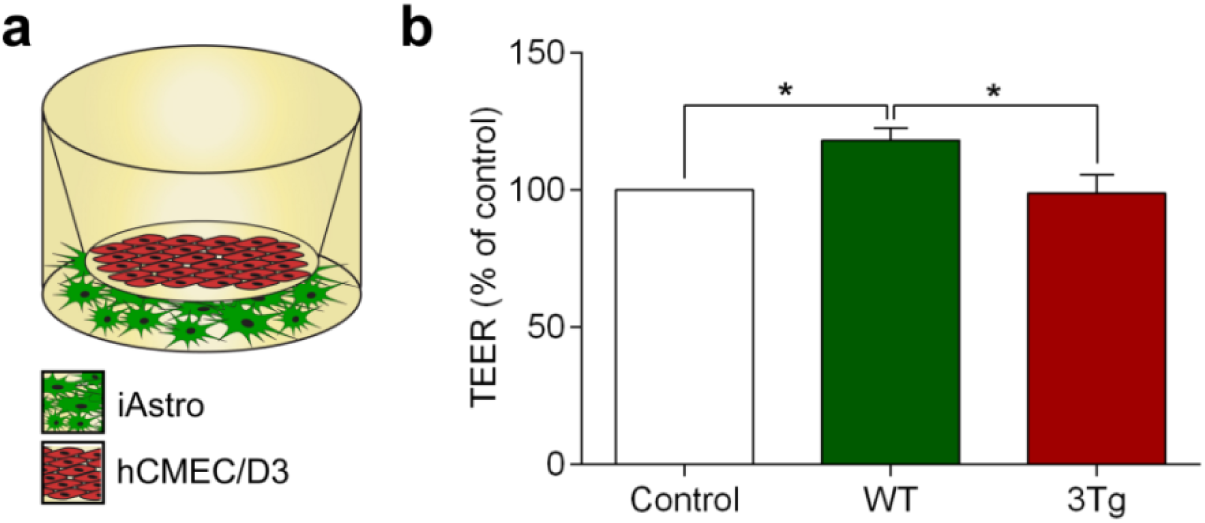
Co-culture with WT-iAstro, but not with 3Tg-iAstro increase transendothelial electrical resistance (TEER) of hCMEC/D3 brain endothelial cell monolayers. (**a**) Schematic representation of the experimental set-up. (**b**) TEER was measured at day 6 of co-culture of hCMEC/D3 monolayers with WT-iAstro, or 3Tg-iAstro. Data represent TEER as a percentage (± SEM) relative to hCMEC/D3 monoculture controls. Statistically significant difference was determined by one way ANOVA Tukey’s test, *p < 0.05 (n = 11-14).

### WT-iAstro but not 3Tg-iAstro up-regulate expression of tight junction proteins in endothelial cells

Paracellular permeability of hCMEC/D3 monolayers is controlled by tight junctions (TJ) composed from specific proteins, which include claudin-5, occludin and Zonula occludens-1 (ZO-1) (Weksler et al. 2013). We hypothesised that in the healthy brain astrocytes secrete certain factors, which up-regulate the expression of endothelial TJ proteins; increased expression of the latter is reflected by an increase in BBB resistance. We therefore tested how co-culturing endothelial cells with WT-iAstro and 3Tg-iAstro affects expression of claudin-5, occludin and ZO-1 in hCMEC/D3s monolayers. Endothelial monolayers were fixed on the polyester membranes the day after TEER measurements (on day 7th of co-culturing with either WT-iAstro or 3Tg-iAstro). Expression of all three proteins was assessed by immunocytochemistry (Fig. 2a) and quantified by Western blotting (Fig. 2b). We found that the expression of all three junctional proteins in hCMEC/D3 cells was significantly higher when co-culturing with WT-iAstro compared to 3Tg-iAstro (in immunocytochemistry the increase was by 106.87 % for claudin-5, n = 5, p < 0.0001; 40.5 % for occludin, n = 5, p < 0.01; and 59.42 % of ZO-1, n = 5, p < 0.05; Fig. 2c). Alternatively, in Westerrn blots all junctional proteins decrease when co-culturing with 3Tg-iAstro compared to control or WT-iAstro (by 36.47 % for claudin-5, n = 4, p < 0.05; 49.22 % for occludin, n = 3, p < 0.05; and 37.27 % of ZO-1, n = 4, p < 0.01; Fig. 2d)

**Fig. 2.**
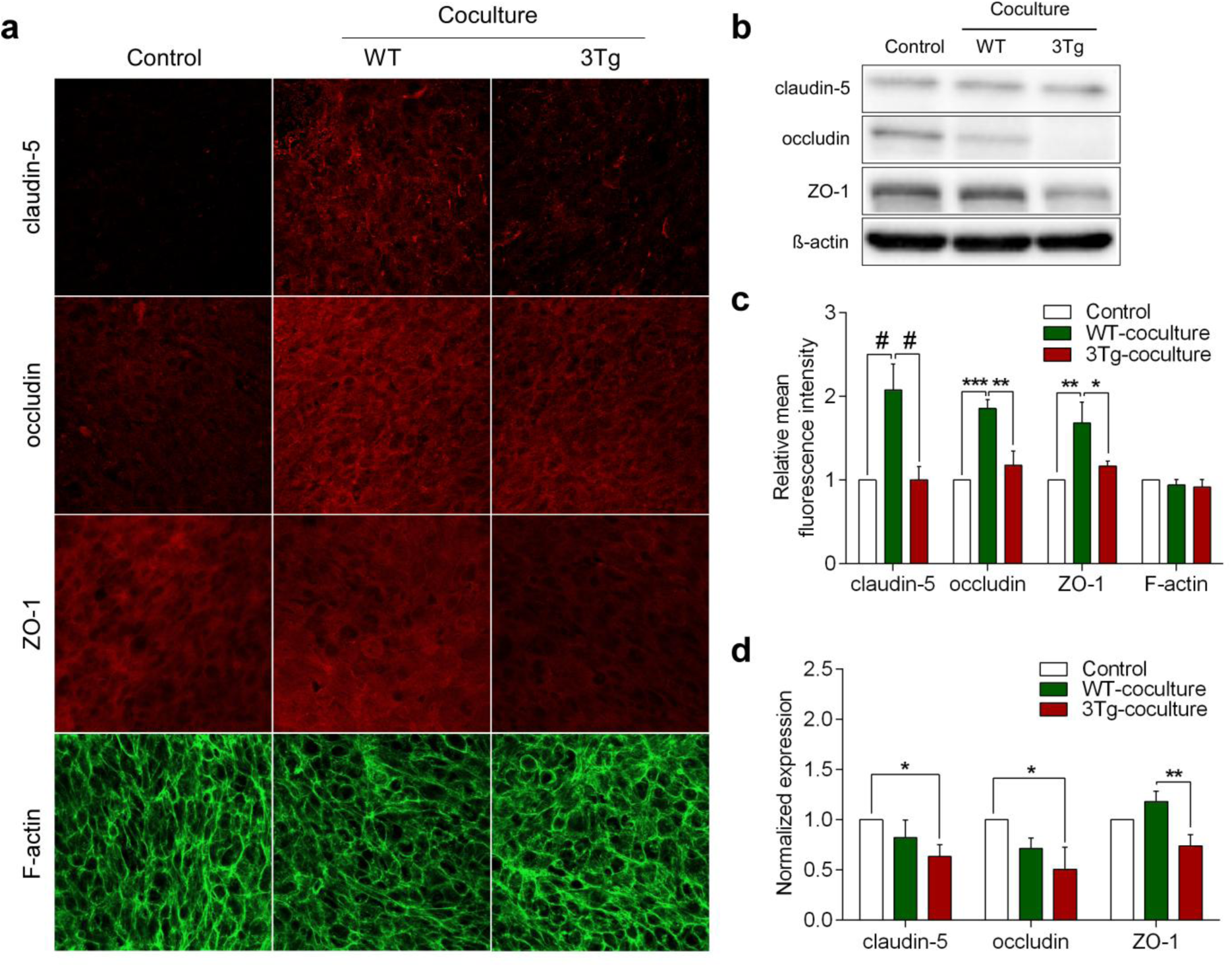
The effects of WT-iAstro and 3Tg-iAstro on the expression of tight junction proteins in hCMEC/D3 monolayers. (**a**) Representative confocal images of endothelial monolayers after 7 days of co-culturing with WT-iAstro, or 3Tg-iAstro; Endothelial monolayers were stained with antibodies against TJ proteins (claudin-5, occludin and ZO1) and F-actin. (**b**) Representative Western blots of tight junction proteins in the endothelial cultures. Full-length Western blot images are provided in supplements; (**c**) Quantification of the mean fluorescence intensity of tight junction proteins in a frame was processed by Leica Application Suite X (LAS X) software (Leica Microsystems). Data represent mean fluorescence intensity (± SEM) relative to hCMEC/D3 monoculture controls. Statistically significant difference was determined by two way ANOVA uncorrected Fisher’s LSD test,*p < 0.05, **p < 0.01, ***p < 0.001, #p < 0.0001 (n = 5). (**d**) Quantification of band intensity of tight junction proteins was performed by Image Lab software. The amount of protein in each lane was determined by ratio of tight junction protein (claudin-5, occludin and ZO1) to β-actin band intensity. Each graph represents mean protein level (± SEM) relative to hCMEC/D3 monoculture controls. Statistically significant difference was determined by two way ANOVA Fisher’s LSD test,*p < 0.05, **p < 0.01 (n = 3-4).

### Extracellular vesicles from WT-iAstro, but not 3Tg-iAstro increase TEER and up-regulate expression of tight junction proteins in hCMEC/D3 cells

Extracellular vesicles (EVs) mediate intercellular communications in various contexts, tissues and organs, including nervous system. Several studies demonstrated that astrocyte-derived EVs from AD patients are enriched in pro-inflammatory cytokines and chemokines indicating possible contribution to the disease (Delpech et al. 2019). We thus isolated EVs from WT-iAstro and 3Tg-iAstro using standard differential centrifugation technique (see Methods). The majority of EVs from both iAstro populations were approximately 140 nm in diameter (Fig. 3a) and expressed classical EV markers (CD63, MFG-E8, Syntenin-1, HSC70; Fig. 3b). Transmission electron microscopy revealed cup-shaped particles, which morphologically corresponded to the EVs (Fig. 3c).

**Fig. 3.**
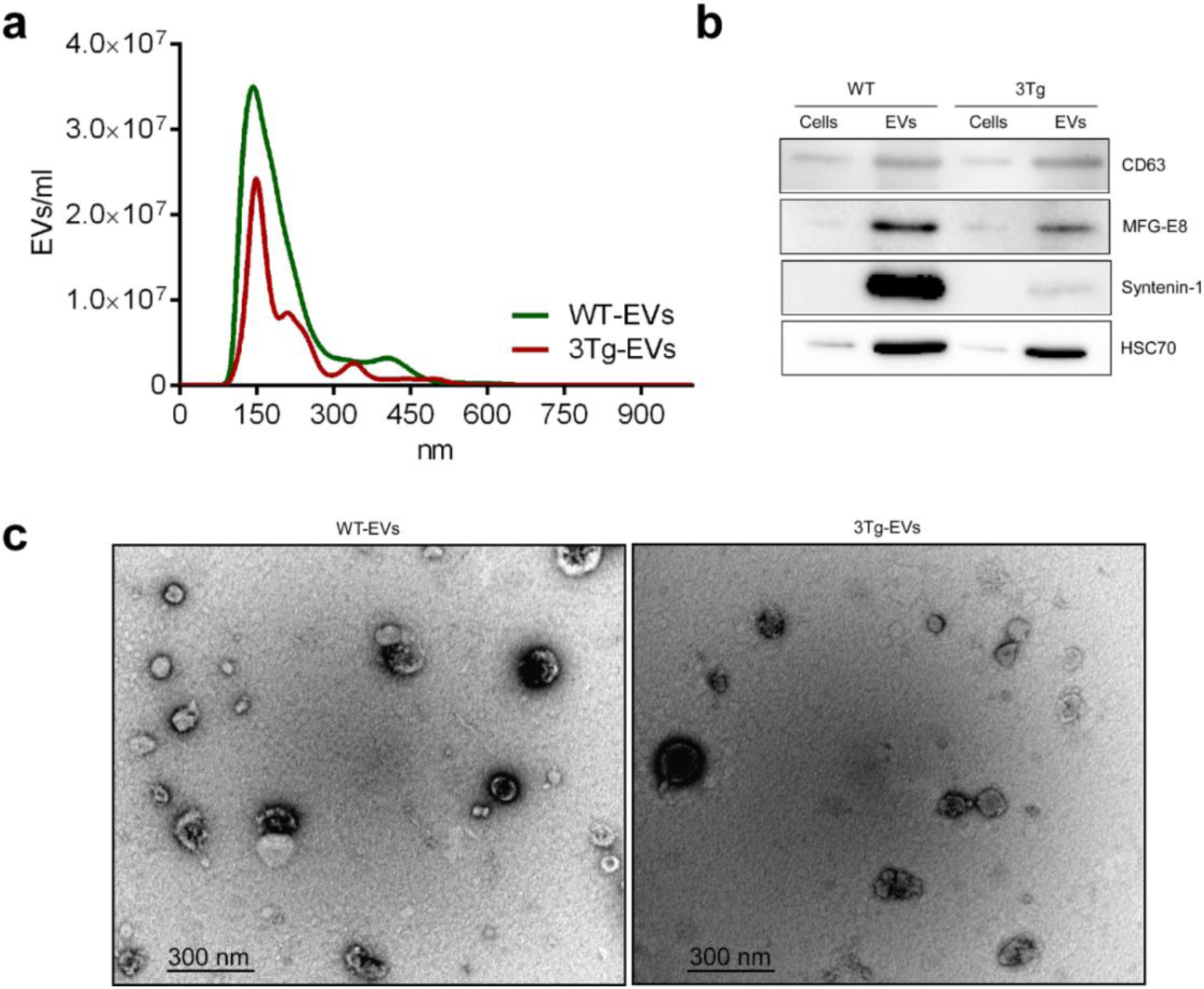
Characterization of extracellular vesicles isolated from WT-iAstro and 3Tg-iAstro. (**a**) iAstro derived EVs Nanoparticle Tracking Analysis (NTA) was performed by NanoSight LM10 instrument (Malvern Panalytical). Size distribution of both WT-iAstro and 3Tg-iAstro derived EVs was around 140 nm (n = 3). (**b**) iAstro cell and EV lysates were subjected to electrophoresis, blotted and the membrane was probed with antibodies against EV markers (CD63, HSP70, MFG-E8, syntenin-1). Bands were visualized by incubation with appropriate horseradish peroxidase-conjugated secondary antibodies and chemiluminescence substrate (n = 3). Unprocessed Western blot images are provided in supplements (**c**) Representative electron microphotographs of EVs derived from WT-iAstro and 3Tg-iAstro. Images were acquired using FEI Morgagn 268 transmission electron microscope, iTEM 5.0 software.

We then compared the effects of EVs derived from WT-iAstro and 3Tg-iAstro on TEER of hCMEC/D3s monolayers. hCMEC/D3 were exposed to iAstro-derived EVs (1.83×10^8^ EVs/insert) for 4 days (Fig. 4a). Incubation of endothelial monolayers with EVs from WT-iAstro increased TEER from 14.11 ± 0.676 W (n = 8) to 17.67 ± 0.213 W or by 25.3 ± 1.51 % (n = 9), p < 0.0001 (Fig. 4b). By contrast, incubation of endothelial monolayers with EVs from 3Tg-iAstro did not significantly affect TEER values in untreated hCMEC/D3 monolayers; the average TEER reading was 14.89 ± 0.413 W (n = 9); this corresponded to a non-significant increase by 5.5 ± 2.92 %, p = 0.2433 (Fig 4b). Changes in TEER reflected changes in expression of TJ proteins: incubation of endothelial monolayers with EVs from WT-iAstro significantly increased expression of tight junction proteins. Immunocystochemistry (Fig. 5a,c) revealed a increase in fluorescence for claudin-5 and ZO-1 by 33.85 %, n = 4, p < 0.05 and by 29.28 %, n = 4, p < 0.01 compared to EVs from 3Tg-iAstro. In western blots expression of claudin-5 and ZO-1 decreased after incubation with EVs from 3Tg-iAstro compared to control or WT-EVs group (by 59.12 %, n = 3, p < 0.001 and by 59.87 %, n = 3, p < 0.05 respectively); changes in expression of occludin were insignificant (Fig 5b,d).

**Fig. 4.**
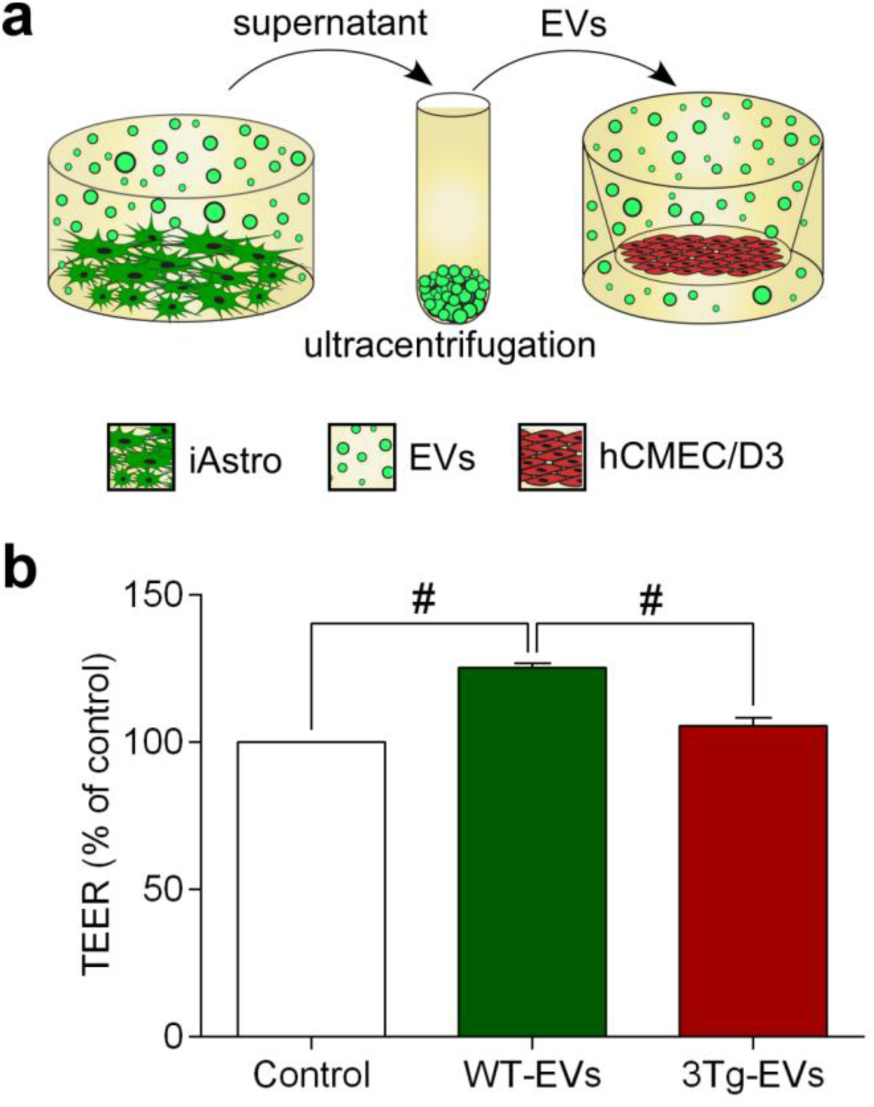
Extracellular vesicles from WT-iAstro, but not 3Tg-iAstro increase TEER of hCMEC/D3 brain endothelial cells. (**a**) Schematic representation of the experimental set-up. (**b**) TEER was measured after 4 days incubation with EVs. Data represent TEER as a percentage (± SEM) relative to untreated hCMEC/D3 controls. Statistically significant difference was determined by one way ANOVA Tukey’s test, #p < 0.0001 (n = 8 -9).

**Fig. 5.**
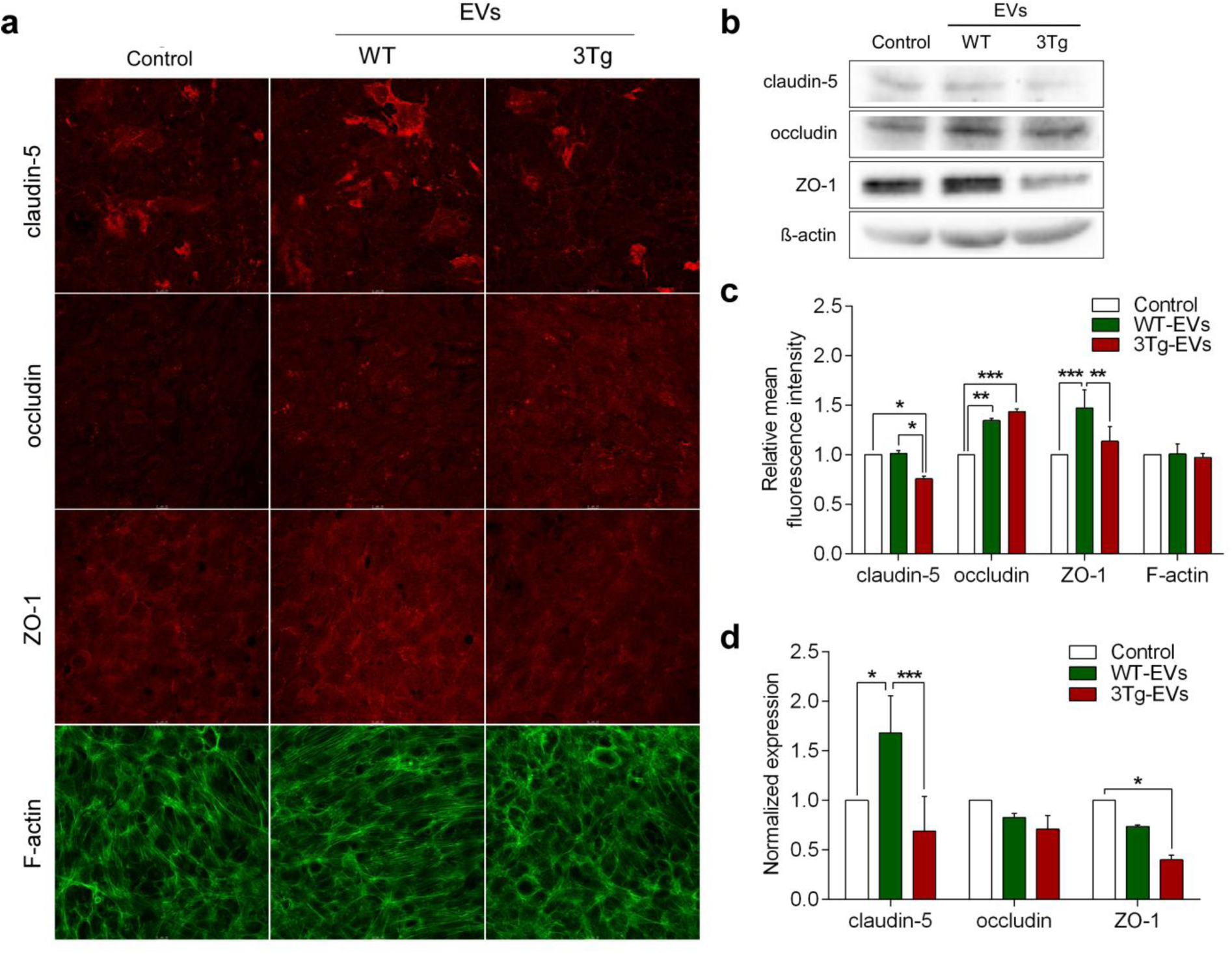
The effects of extracellular vesicles from WT-iAstro and 3Tg-iAstro on the expression of tight junction proteins in hCMEC/D3 monolayers. (**a**) Representative confocal images of endothelial monolayers staind with antibodies against tight junction proteins (claudin-5, occludin and ZO1) and F-actin. (**b**) Representative Western blots of tight junction proteins and b-actin. Full-length Western blot images are provided in supplements. (**c**) Quantification of the mean fluorescence intensity of tight junction proteins in a frame was processed by Leica Application Suite X (LAS X) software (Leica Microsystems). Data represent mean fluorescence intensity (± SEM) relative to untreated hCMEC/D3 controls. Statistically significant difference was determined by two way ANOVA uncorrected Fisher’s LSD test,*p < 0.05, **p < 0.01, ***p < 0.001 (n = 4). (**d**) Quantification of band intensity of tight junction proteins was performed by Image Lab software. Each graph represents mean protein level (± SEM) determined by ratio of tight junction protein (claudin-5, occludin and ZO1) to β-actin band intensity relative to untreated hCMEC/D3 controls. Statistically significant difference was determined by two way ANOVA Fisher’s LSD test,*p < 0.05, ***p < 0.001 (n = 3).

### Endothelial cells to astrocytes communication distinctly regulate expression of selected genes in WT-iAstro and 3Tg-iAstro cells

In the operational BBB endothelial cells and astrocytes are functionally connected through bi-directional communications mediated by various secreted factors. To test for endothelial cells to astrocytes communication we analysed how conditioned medium from hCMEC/D3s affects expression of selected genes in WT-iAstro and 3Tg-iAstro. For this purpose iAstro were incubated in the conditioned medium (CM) from hCMEC/D3s for 48 hours (with one medium change after 24 hours, Fig. 6a). We found that treatment with CM from hCMEC/D3s increased expression of pro-inflammatory chemokine CCL2 in both WT-iAstro and 3Tg-iAstro. Elevation of the expression of CCL2 in 3Tg-iAstro + CM group was significantly higher compared to WT-iAstro + CM group (10.47 ± 1.28, n = 3 for 3Tg-iAstro + CM and 3.57 ± 0.96, n = 3 for WT-iAstro + CM, p = 0.0006; Fig. 6b). When compared to CM-treated WT-iAstro, 3Tg-iAstro also showed increased expression of pro-inflammatory cytokine IL6 (3.05 ± 0.32, n = 3 in 3Tg-iAstro + CM and 1.27 ± 0.3, n = 3 in WT-iAstro + CM, p = 0.0032; Fig. 6b). Subsequently performed ELISA confirmed qPCR data and revealed significant increase of CCL2 (by 7,83 ± 2.75 folds, n = 4 compared to control, p < 0.05) but not IL6 (by 2.32 ± 0.75 folds n = 3 compared to control) presence in the supernatants from 3Tg-iAstro cultures incubated in CM from hCMEC/D3 (Fig. 6c).

**Fig. 6.**
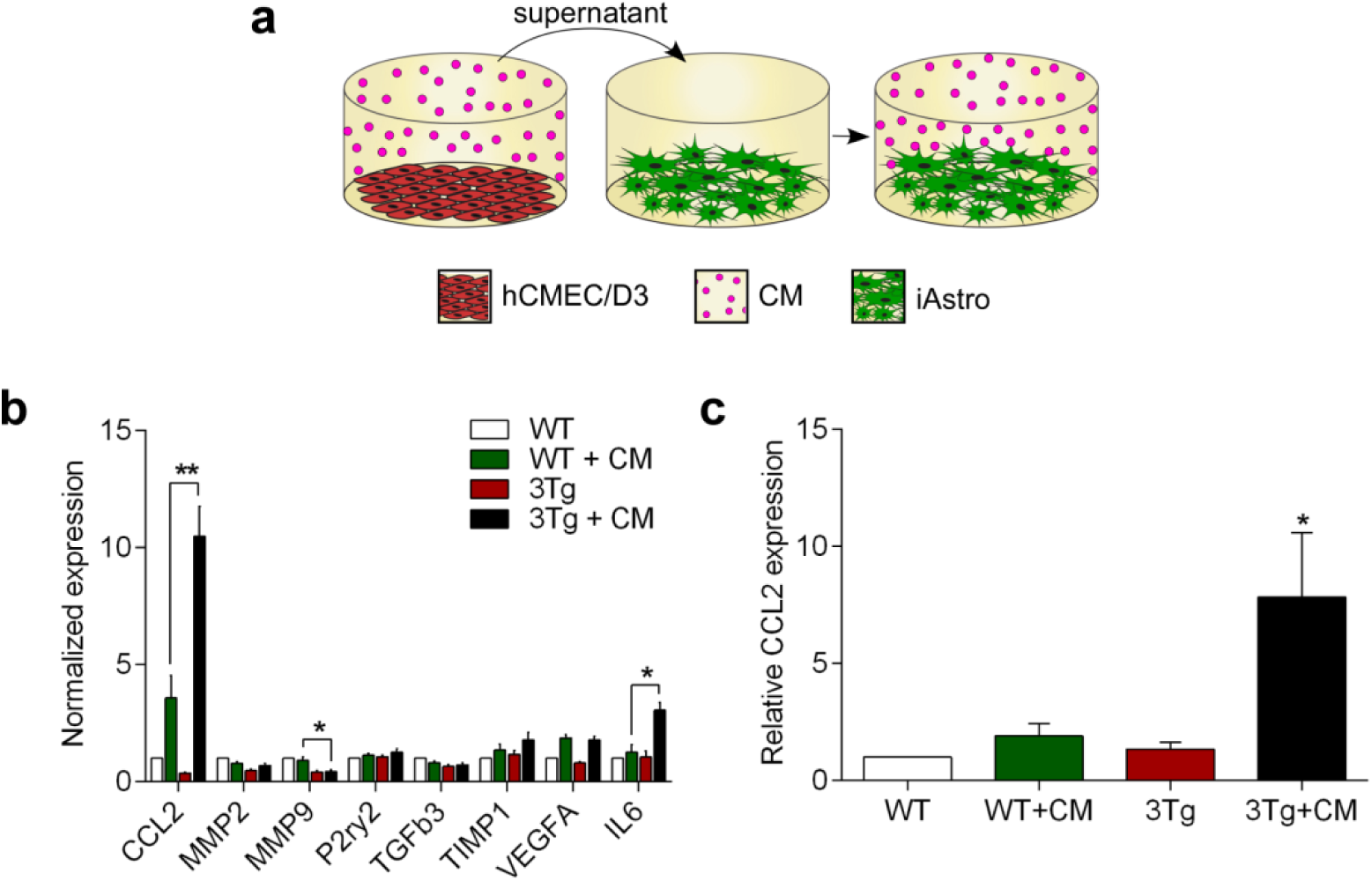
Conditioned medium from hCMEC/D3 brain endothelial cells increase expression and secretion of pro-inflammatory chemokine CCL2 in 3Tg-iAstro cells. (**a**) Schematic representation of the experimental set-up. (**b**) Pre-conditioned (for 24 hours) EndoGRO medium from hCMEC/D3s was administered to WT-iAstro and 3Tg-iAstro cells (previously cultured in EndoGRO) for 24 hours; subsequently cells were treated with fresh pre-conditioned medium for another 24 hours and total RNA was isolated for gene expression analysis. qPCR reactions were performed in duplicates and experiments repeated at least three times. The expression of genes was quantified using the CFX 96 instrument and data analysed by CFX Manager 3.1 software, *p < 0.01, **p < 0.001 (n = 3). (**c**) The levels of CCL2 secretion in WT-iAstro and 3Tg-iAstro cells were evaluated after 24 hour incubation with EndoGRO (control), or after exposure to the pre-conditioned (for 24 hours) EndoGRO medium (CM) from hCMEC/D3s. The levels of CCL2 were quantified with mouse CCL2 ELISA kit (Thermo Fisher Scientific, Cat. Nr. BMS6005) according to the manufacturer’s instructions. Data represent mean (± SEM) relative to WT-iAstro. Statistical significance was determined by one-way ANOVA Tukey’s test, *p > 0.05 (n = 4).

## Discussion

Neurovascular dysfunction affects all components of neurovascular unit (NVU) and contributes to AD pathogenesis from the earliest stages (Kisler et al. 2017; Sweeney et al. 2018). Astrocytes support BBB integrity through close interactions with other components of the NVU. In addition to providing structural support by formation of perivascular endfeet and basal membrane (glia limitans) astrocytes secrete numerous paracrine factors that have been implicated in the complex and multilayered astrocyte-endothelial (Alvarez et al. 2011) and astrocyte-pericyte (Bell et al. 2012) crosstalk to regulate barrier properties of the BBB. In particular astrocytes are known to secrete sonic hedgehog and retinoic acid which up-regulate expression of TJ proteins in endothelial cells (Alvarez et al. 2011).

However, the role of astrocytes during BBB breakdown occurring at the early stages of AD remains unclear. In the present study by using membrane-based *in vitro* BBB model we confirmed the fundamental role of astrocytes in supporting BBB integrity. In the chimeric BBB model composed of human hCMEC/D3 cells and WT astrocytes we found (i) significant up-regulation of TJ proteins expression by endothelial cells and (ii) an increase in electrical resistance of endothelial monolayer. Immortalised astrocytes obtained from 3xTg-AD mice failed to produce such changes, confirming that these cells retain pathological signature associated with the disease. We used cell contact-free co-culture model, therefore the observed effects were due to the factors secreted by astrocytes. Astroglia-derived paracrine factors are essential for the formation and maintenance of BBB (Sweeney et al. 2019; Alvarez et al. 2013), thus impaired capability of 3Tg-iAstro to support BBB may reflect deficient secretion of such paracrine factor(s). Proteomic comparison of supernatants from endothelial/iAstro co-cultures is needed to support, or reject this possibility.

Another pathway of astroglia-endothelial communication may be mediated through EVs. It is now widely accepted that composition and biological properties of EVs largely depends on the parent cell type and its physiological or pathological state. Thus, EVs may act as promoters, or suppressors of pathological processes in CNS (Delpech et al. 2019). Astrocyte-derived EVs have been implicated into the cell-to-cell transmission of phospho-Tau protein (Chiarini et al. 2017). Astrocyte-derived exosomes from plasma samples of AD patients contain high levels of complement proteins and pro-inflammatory cytokines (Goetzl et al. 2018). Here we demonstrate that EVs derived from WT-iAstro and 3Tg-iAstro exhibit the same effects on the BBB integrity as their parent cells. Different functional properties of EVs from WT-iAstro and 3Tg-iAstro imply significant differences in their cargo content. Comparative proteomic and RNAomic analyses are needed in order to identify proteins and (or) microRNAs responsible for the observed effects. It is presently unclear, whether the effects observed in co-culture experiments were also EV-dependent, or they were caused by other paracrine factors secreted by astrocytes. It should be noted, that according to our estimations EV doses used in our experimental setting were by several times higher than those produced by astrocytes during co-culture with hCMEC/D3s.

Alternatively, 3Tg-iAstro could secrete factors negatively affecting BBB integrity. Indeed, astrocytes secrete numerous pro-inflammatory factors which may disrupt the BBB and promote leukocyte migration into the CNS (Sofroniew 2015). For example, several studies reported that pro-inflammatory chemokine CCL2 (MCP-1) may alter the endothelial tight junction (TJ) structure and compromise BBB integrity. The CCL2 caused reorganization of actin cytoskeleton with stress fiber formation in brain endothelial cells (Stamatovic et al. 2003). Treatment with CCL2 increased phosphorylation of adaptor proteins ezrin, radixin and moesin which then bind to the protein ZO-1 and pull it away from the TJ, leading to BBB deficiency (Yao et al. 2011). Also, CCL2 induced phosphorylation and redistribution of TJ proteins ZO-1, ZO-2, occludin and claudin-5 *via* RhoA, ROCK and PKCα signalling pathways (Stamatovic et al. 2006). We found that treatment with conditioned medium from hCMEC/D3s significantly increased secretion of CCL2 (MCP-1) in 3Tg-iAstro, but not in WT-iAstro. We therefore suggest that increased secretion of CCL2 by 3Tg-iAstro may contribute to TEER decrease in for hCMEC/D3s monolayers.

In conclusion, we show for the first time that immortalised hippocampal astrocytes from 3xTG-AD mice exhibit impaired capacity to support BBB integrity *in vitro* through paracrine mechanisms. The peculiarity of our co-culture model is the employments of cross-species cells as well as the usage of immortalized cell lines. This model however demonstrated that even after immortalization astrocytes from 3xTG mice remain pathologically remodeled. This finding further extends the concept of astroglial atrophy and loss of function, as indeed astroglial inability to support BBB integrity may represent an important factor underlying vascular abnormalities contributing to the pathophysiology of AD.

## Methods

### Cell lines

#### Immortalized astroglial cell lines

Immortalized astroglial cell lines used in this study were provided by Dr. Dmitry Lim (Rocchio et al. 2019). These lines have been established from the hippocampi of 3xTg-AD mice model carrying PS1(M146), APP(Swe) and Tau(P301L) mutations (3xTg-iAstro), or from wild-type control mice (WT-iAstro). Both cell lines were maintained in a plastic cultureware in low glucose (1 g/L) DMEM-GlutaMAX™ medium (Gibco) supplemented with 10 % foetal bovine serum (FBS; Biochrom), 100 U/ml penicillin-streptomycin (Biochrom), hereafter referred to as iAstro medium. WT-iAstro and 3xTg-iAstro cells used for the experiments were between passages 14 and 20.

#### Endothelial hCMEC/D3 cell line

Immortalized human cerebral microvascular endothelial cell line (hCMEC/D3) was purchased from Merck-Millipore (Darmstadt, Germany). All cultureware used for hCMEC/D3 cells was coated with rat tail collagen type I (Gibco) diluted in phosphate buffered saline (PBS) 1:20 and incubated at 37 °C for at least 1 hour. Cells were maintained in EndoGRO-MV complete medium (Merck-Millipore) containing EndoGRO basal medium, 5% FBS, L-glutamine (10 mM), EndoGRO-LS supplement (0.2 %), heparin sulphate (0.75 U/ml), ascorbic acid (50 μg/ml), hydrocortisone hemisuccinate (1 μg/ml), recombinant human epidermal growth factor (5 ng/ml) and additionally supplemented with freshly added recombinant human fibroblast growth factor-basic (1 ng/ml; Gibco). hCMEC/D3 cells used for the experiments were between passages 6 and 11. All cell lines were maintained at 37 °C in a humidified atmosphere containing 5 % CO_2_. The medium was changed twice a week. For passaging, cells were washed once with PBS, harvested with 0.25 % trypsin/ 1 mM EDTA solution (Gibco), resuspended in culture medium and plated onto cell culture flask for expansion.

### Co-culture of hCMEC/D3 and iAstro

iAstro and hCMEC/D3 co-cultures were studied using 0.33 cm^2^ transparent polyester membrane transwell inserts with pore size of 0.4 μm in 24-well plate format (Corning). iAstro lines were plated in a well plate at a seeding density of 2×10^4^ cells/cm^2^ and allowed to attach for 2 hours. hCMEC/D3 cells were then plated on the top side of the collagen pre-coated membrane of transwell insert at a seeding density of 6×10^4^ cells/cm^2^. We used different combinations of hCMEC and iAstro for co-culture experiments: hCMEC/D3-blank control, hCMEC/D3-WT-iAstro and hCMEC/D3-3xTg-iAstro. Alternatively, hCMEC/D3 cells were plated on the top side of the collagen pre-coated transwell insert at a seeding density of 6×10^4^ cells/cm^2^ and allowed to reach confluence. At day 4 after seeding hCMEC/D3s were exposed to complete EndoGRO medium containing iAstro-derived extracellular vesicles (EVs) at a concentration of 1.83×108 EVs per insert for the next 4 days. Complete EndoGRO medium, including medium containing EVs, was used in both compartments of transwell system.

### Transendothelial electrical resistance (TEER)

TEER measurements were performed on day 6th of co-cultivation of iAstro and hCMEC/D3, or on the day 4th after exposure of iAstro-derived EVs on hCMEC/D3s, 1 hour after replacement of complete EndoGRO medium, using a Millicell ERS-2 Electrical Resistance System (Merck-Millipore). To calculate transendothelial electrical resistance (TEER; Ω×cm^2^), the electrical resistance across the insert membrane without cells was subtracted from the readings obtained on inserts with cells, and this value was multiplied by the surface area of the insert (0.33 cm^2^).

### Isolation of extracellular vesicles

EVs from iAstro cells were isolated using differential centrifugation protocol according to the Thery et al. (2018) with some modifications. Briefly, iAstro cells were cultivated in iAstro EV-depleted medium for 3-4 days. Then the supernatants were collected and immediately subjected to several ultracentrifugation steps at 4 °C: 300 g for 10 min, 2000 g for 10 min and 20 000 g for 30 min. Finally supernatants were ultracentrifuged at 100 000 g for 70 min at 4 °C in Sorvall LYNX 6000 ultracentrifuge, with titan rotor T29-8×50 in oak ridge centrifuge tubes with sealing caps (all from Thermo Fisher Scientific, Rochester, NY). After removal of the supernatants, pellets were resuspended in ice-cold PBS and ultracentrifuged again at 100000 g for 70 min at 4 °C using the same tubes and rotor. Final pellets of EVs were resuspended in ice-cold PBS and stored at -80 °C.

Nanoparticle tracking analysis (NTA) was performed with NanoSight LM10 (Malvern Panalytical). Before measurement all EV preparations from individual collections were pooled and diluted 33 times by adding 15 μl of EV pool to 485 μl of PBS. Size distribution of the EVs from both cultures (WT and 3Tg) was around 140 nm. According to the NTA measurements single dose of EV contained 1,83×10^8^ vesicles.

### Transmission electron microscopy

Transmission electron microscopy (TEM) of EVs was performed according to the previously reported protocol ^39^ with some modifications. Briefly, EVs in PBS were fixed in 2 % paraformaldehyde for 40 min on ice. Formvar-carbon coated copper grids were floated on a 10 µl drops of fixed EV suspension for 20 min at room temperature. Then the grids were washed with PBS and floated on a 30 µl drops of 1% glutaraldehyde for 5 min at room temperature, then again washed eight times by transferring from one drop of distilled water to another. Samples were contrasted on 30 µl drops of 2 % neutral uranyl acetate for 5 min at room temperature in the dark. Finally, grids were air dried for 5 min. All incubations were displayed on a Parafilm sheet with the coated sides of grids facing to the drop. The samples were analysed with the transmission electron microscope FEI Morgagni 268.

### RNA extraction, cDNA synthesis and real-time PCR

Pre-conditioned (24 hours) EndoGRO medium from hCMEC/D3s was subjected onto WT-iAstro and 3Tg-iAstro cells (previously cultivated in EndoGRO). After 24 hours incubation WT-iAstro and 3Tg-iAstro cells were treated again with fresh pre-conditioned medium for another 24 hours. Total RNA was isolated using miRNeasy RNA purification kit (Qiagen) according to the manufacturer’s instructions. Genes RNA expression assessed by RT-qPCR method. cDNA synthesized using Maxima First strand cDNA synthesis Kit (Thermo Scientific) and for quantitative PCR (qPCR) was used Maxima SYBR Green qPCR master mix (Thermo Scientific), according to the manufacturer’s instructions. Briefly, 1 μg of isolated RNA converted into cDNA in 35 μl reaction mixture. qPCR performed using 0.25 μl of synthesised cDNA and 0.2 μM primer pairs in 25 μl reaction volume. PCR cycling was used as follows: initial denaturation 95°C – 10 min, 40 cycles of 95°C – 10 s, 55°C – 30 s and 72°C – 30 s. Specificity of the product assessed by melting curve analysis. Relative gene expression analysed using 2-ΔΔCT method. As reference gene was used hypoxanthine guanine phosphoribosyl transferase (Hprt). All reactions performed using the CFX 96 instrument, data analysed by CFX Manager 3.1 software (Bio-Rad, Hercules, CA).

### Protein isolation and Western blot analysis

After 7 days in co-culture with iAstro (in transwell inserts), or at day 4 after exposure to EVs (in 24-well plate), hCMEC/D3s were washed three times with room temperature (RT) PBS and lysed using Pierce RIPA buffer (Thermo Scientific) containing 1x Halt protease inhibitor cocktail (Thermo Scientific) for 15 min on ice. Lysates were collected, vortexed and placed on ice again for the next 10 min. Afterwards lysates were centrifuged at 18 000 g for 20 min at 4 °C to remove cell debris. Protein concentration of remaining supernatants was measured with the NanoPhotometer Pearl (Implen). For Western blot analysis cell lysates were diluted in 6x Laemmli sample buffer and denatured for 5 min at 95 °C. Equal amount of proteins from each lysate were subjected to polyacrylamide gel (4-10 %) electrophoresis (PAGE) in Mini-PROTEAN Tetra cell apparatus (Bio-Rad). PAGE separated proteins were blotted onto a PVDF membrane in a semi-dry Trans-Blot Turbo transfer system (Bio-Rad). The PVDF membrane was blocked with 5 % bovine serum albumin (BSA; Applichem)/PBS-Tween for 1 hour RT on a platform rocker. The membrane was then probed with the following primary antibodies diluted in 5 % BSA /PBS-T for overnight at 4 °C: β-actin (1:1000; Cell Signaling Technology), ZO-1 (1.25 μg/ml), claudin-5 (1 μg/ml) and occludin (2 μg/ml; all from Thermo Fisher Scientific). After incubation with primary antibodies membrane was washed with PBS-T three times on a platform rocker and incubated with horseradish peroxidase-conjugated secondary antibody diluted in PBS-T 1:2000 (Thermo Scientific) for 1 hour at RT on a platform rocker. Washing procedure was repeated and immunoreactive bands were detected with Clarity ECL Western blotting substrate (Bio-Rad) using ChemiDoc MP system (Bio-Rad).

### Confocal microscopy

After 7 days of co-culture with iAstro (in transwell inserts), or at day 4 after exposure of EVs (on cover slips), hCMEC/D3 were fixed with 4% paraformaldehyde (PFA) for 20 min, washed with PBS three times, permeabilized with 0.1% Triton X-100 for 15 min and washed with PBS three times again. Afterwards, the samples were blocked with 1 % BSA for 30 min (all procedures were performed at RT). After blocking, cells were incubated overnight at 4 °C with primary antibody against ZO-1 (7.5 μg/ml, Thermo Fisher Scientific). Alternatively, cells were fixed and permeabilized with 100 % methanol for 20 min at -20 °C, then blocked and probed with the following primary antibodies: claudin-5 (1:100) and occludin (1:50), both from Thermo Fisher Scientific). All primary antibodies were diluted in 1% BSA. After incubation with primary antibodies, cells were washed with PBS and incubated with Alexa Fluor 594-conjugated secondary antibodies (1:1000; Life Technologies) or Alexa Fluor 647-phalloidin conjugate (for primary antibody free samples; 1:40; Life Technologies) for 1 hour at RT in the dark. Washing procedure was repeated and cells were coversliped using an aqueous fluorescent mounting medium (Dako). The specimens were analysed under a confocal microscope Leica TCS SP8 (Leica Microsystems) using a DPSS 561 nm and a HeNe 633 nm lasers. Images were taken with a 63x oil immersion objective. Quantification of the mean fluorescence intensity in a frame was processed by Leica Application Suite X (LAS X) software (Leica Microsystems).

### ELISA

The levels of CCL2 secretion in WT-iAstro and 3Tg-iAstro cells were evaluated after 24 hour incubation with EndoGRO (control), or after exposure to the pre-conditioned (for 24 hours) EndoGRO medium from hCMEC/D3s. Cell culture supernatants were centrifuged at 18 000 g for 5 min and stored - 20° C until further analysis. The levels of CCL2 (MCP-1) were quantified with mouse CCL2 ELISA kit (Thermo Fisher Scientific, Cat. Nr. BMS6005) according to the manufacturer’s instructions. The colorimetric measurements (λ=450 nm, reference λ=620 nm) were performed using Asys UVM340 plate reader (Biochrom).

### Statistics

Statistical analysis was performed from at least three biological experiments with at least two technical replicates in each if not indicated differently. Data represent mean and standard error of the mean (SEM) values. Difference between three or more groups were compared by one-way ANOVA following uncorrected Tukey’s post-test. Grouped data were analysed by two-way ANOVA followed by uncorrected Fisher’s LSD post-test. Graph Pad Prism® software version 6.0 (Graph Pad Software, Inc., USA) was used for all data analysis if not indicated differently. Real-time PCR experiments were performed in triplicates and SEM values were calculated, a statistical analysis of the data was performed using a one-way ANOVA followed by a Dunnett’s test using MaxStat Pro Statistics Software (Version 3.6).

## Acknowledgments

We are grateful to Dr. Aistė Jekabsonė for help with cell lines.

