## Supplemental Figure 1 for "Immortalised hippocampal astrocytes from 3xTG-AD mice fail to support BBB integrity *in vitro*: Role of extracellular vesicles in glial-endothelial communication"

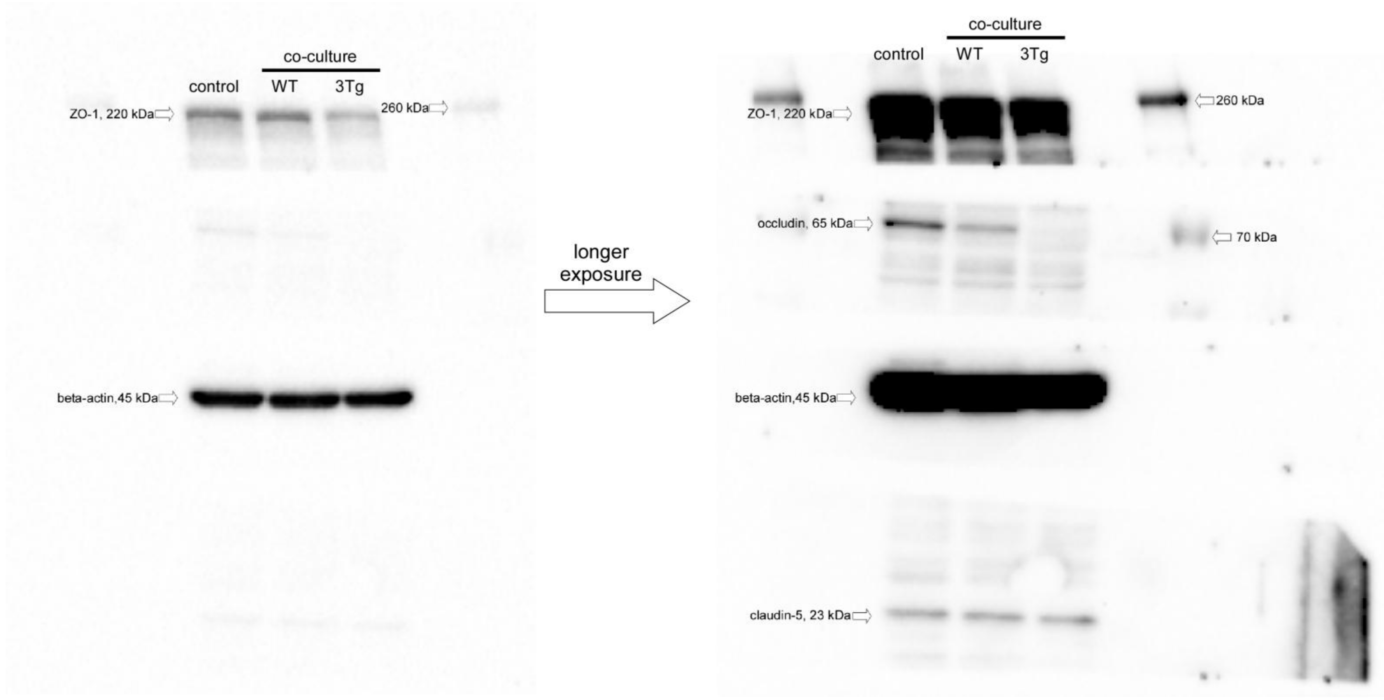


**Supplementary figure 1. Representative unprossesed Western blots of tight junction proteins in the endothelial cells after 7 days of co-culturing with WT-iAstro, or 3Tg-iAstro**. hCMEC/D3 cells were subjected to electrophoresis, blotted and the membrane was probed with antibodies against tight junction proteins (ZO-1, occludin, beta-actin and claudin-5). Bands were visualized by incubation with appropriate horseradish peroxidase-conjugated secondary antibodies and chemiluminescence substrate. Bands were detected by Image Lab software at different exposure times. For quantification were used bands detected before saturation appears. Controls represent hCMEC/D3 monoculture, WT - hCMEC/D3 co-cultured with WT-iAstro, 3Tg - hCMEC/D3 co-cultured with 3Tg-iAstro.
