## Supplemental Figure 2 for "Immortalised hippocampal astrocytes from 3xTG-AD mice fail to support BBB integrity *in vitro*: Role of extracellular vesicles in glial-endothelial communication"

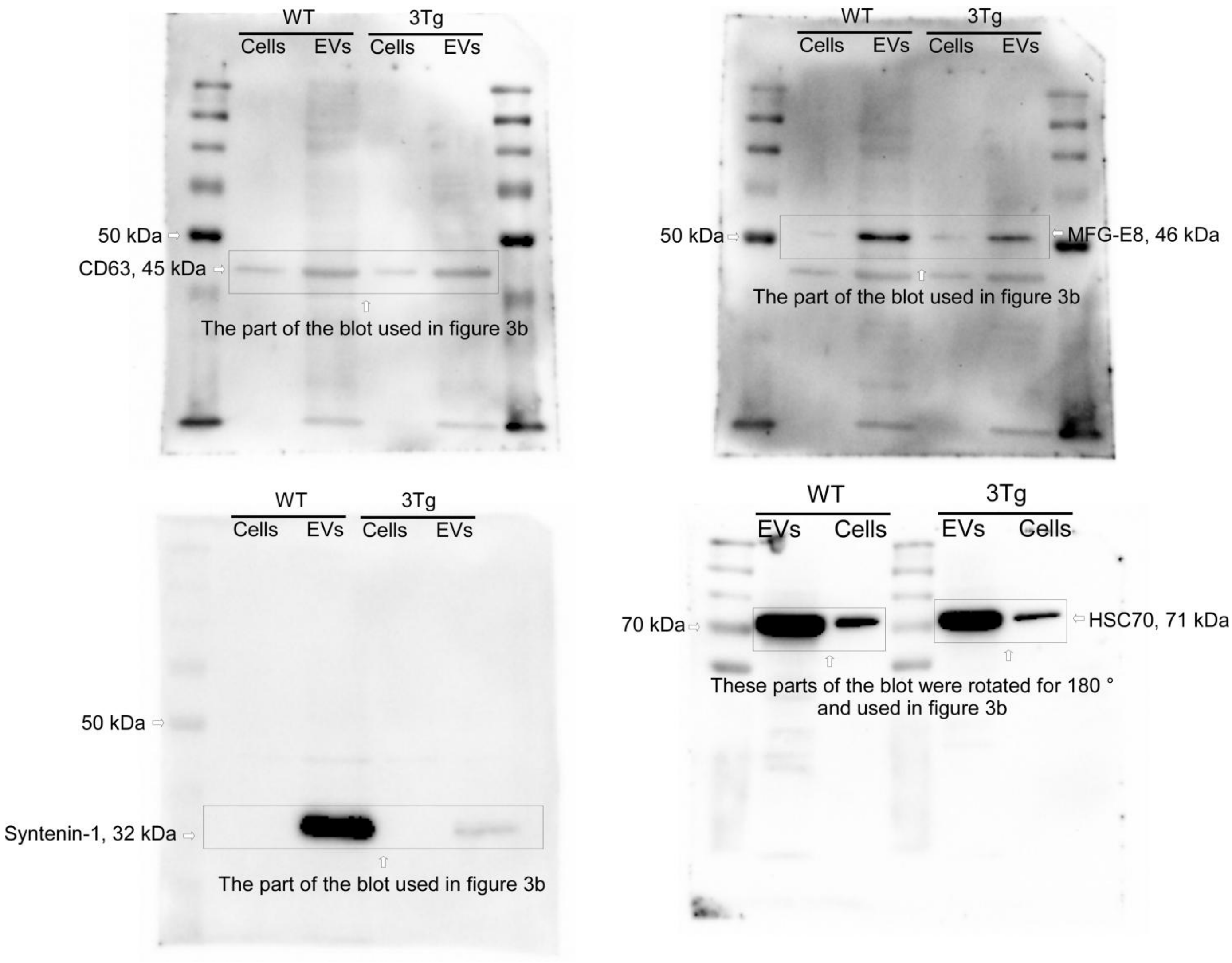


**Supplementary figure 2. Representative unprossesed full-lenght Western blots of extracellular vesicles isolated from WT-iAstro and 3Tg-iAstro**. iAstro cell lines and EV lysates were subjected to electrophoresis, blotted and the membrane was probed with antibodies against EV markers (CD63, MFG-E8, syntenin-1, HSP70). Bands were visualized by incubation with appropriate horseradish peroxidase-conjugated secondary antibodies and chemiluminescence substrate. Bands were detected by Image Lab software at different exposure times.
